# Transcriptional profiling of isogenic Friedreich ataxia induced pluripotent stem cell-derived neurons

**DOI:** 10.1101/457093

**Authors:** Jiun-I Lai, Daniel Nachun, Lina Petrosyan, Benjamin Throesch, Erica Campau, Fuying Gao, Kristin K. Baldwin, Giovanni Coppola, Joel M. Gottesfeld, Elisabetta Soragni

**Affiliations:** Department of Molecular Medicine, La Jolla, CA, USA; Department of Neuroscience, The Scripps Research Institute, La Jolla, CA, USA; Semel Institute for Neuroscience & Human Behavior, UCLA, Los Angeles, CA, USA

**Author notes:** Correspondence: Elisabetta Soragni, Department of Molecular Medicine, The Scripps Research Institute, 10550 North Torrey Pines Road, La Jolla, CA 92037, USA. These authors contributed equally to this work (Jiun-I Lai, Daniel Nachun). Present address: Jiun-I Lai, Institute of Clinical Medicine, National Yang-Ming University, Taipei, Taiwan.

## Abstract

Friedreich ataxia (FRDA) is a rare childhood neurodegenerative disorder with no effective treatment. FRDA is caused by transcriptional silencing of the *FXN* gene and consequent loss of the essential mitochondrial protein frataxin. Based on the knowledge that a GAA•TTC repeat expansion in the first intron of *FXN* leads to heterochromatin formation and gene silencing, we have shown that members of the 2-aminobenzamide family of histone deacetylase inhibitors (HDACi) reproducibly increase *FXN* mRNA levels in induced pluripotent stem cell (iPSC)-derived FRDA neuronal cells and in peripheral blood mononuclear cells from patients treated with the drug in a phase I clinical trial. How the reduced expression of frataxin leads to neurological and other systemic symptoms in FRDA patients remains unclear. Similarly to other triplet repeat disorders, it is not known why only specific cells types are affected in the disease, primarily the large sensory neurons of the dorsal root ganglia and cardiomyocytes. The combination of iPSC technology and genome editing techniques offers the unique possibility of addressing these questions in a relevant cell model of the disease, without the confounding effect of different genetic backgrounds. We derived a set of isogenic iPSC lines that differ only in the length of the GAA•TTC repeats, using “scarless” gene-editing methods (helper-dependent adenovirus-mediated homologous recombination). To uncover the gene expression signature due to GAA•TTC repeat expansion in FRDA neuronal cells and the effect of HDACi on these changes, we performed transcriptomic analysis of iPSC-derived central nervous system (CNS) and isogenic sensory neurons by RNA sequencing. We find that multiple cellular pathways are commonly affected by the loss of frataxin in CNS and peripheral nervous system neurons and these changes are partially restored by HDACi treatment.

## Introduction

Friedreich ataxia (FRDA; OMIM 22930) is an autosomal recessive neurodegenerative disorder, characterized by progressive spinocerebellar ataxia, dysarthria, sensory neuropathy and pyramidal weakness (1). Most patients show signs of hypertrophic cardiomyopathy (2), about half present with glucose intolerance and 12% are clinically diagnosed with diabetes (3). The initial site of neurodegeneration is the dorsal root ganglia (DRG), followed by spinocerebellar tracts and the dentate nucleus of the cerebellum (2). The DRG show a reduction in size and number of sensory neurons accompanied by proliferation of satellite cells (4). Larger neurons of the proprioceptive type are mainly affected, resulting in loss of position sense, but pain, light touch and temperature perception can also decrease in advanced stages of the disease (1). Analysis of cell sizes in the DRG are consistent with a pathogenic mechanism of hypoplasia rather than atrophy and this supports the idea that normal development of the DRG and the spinal cord is hindered in FRDA (5).

An expanded GAA•TTC triplet repeat sequence, located in the first intron of the nuclear *FXN* gene, causes inhibition of transcription and the loss of the essential mitochondrial protein frataxin in affected individuals. Available evidence supports a role for frataxin in the biogenesis of iron-sulfur (Fe-S) clusters in mitochondria, resulting in impaired activities of Fe-S enzymes, altered cellular iron metabolism, decreased mitochondrial energy production and increased oxidative stress (6,7). To counteract these abnormalities, antioxidants, iron chelators and stimulants of mitochondrial biogenesis have been proposed as therapeutics (8). However, no clear results supporting benefit of any of these drugs have so far been obtained in randomized human trials (9). Other avenues for therapeutic development, therefore, are being pursued, including strategies aimed at increasing frataxin expression by preventing frataxin degradation (10), repeat-targeted oligonucleotides (11) and synthetic transcription elongation factors (12), together with protein replacement therapy (13), stem cell therapy (14) and gene therapy (15). Based on the knowledge that GAA•TTC expansion leads to heterochromatin formation and gene silencing, we have shown that members of the 2-aminobenzamide family of histone deacetylase inhibitors (HDACi) reproducibly increase *FXN* mRNA levels in FRDA lymphoblast cell lines (16), primary lymphocytes from FRDA patients(17), FRDA mouse models (18,19) and human FRDA neuronal cells derived from patient induced pluripotent stem cells (iPSCs)(20). A phase I clinical trial with HDACi **109** (RG2833) demonstrated increased *FXN* mRNA in peripheral blood mononuclear cells (PBMCs) from patients treated with the drug (20) and a derivative of **109** with improved pharmacological properties has been selected as drug candidate by Biomarin Pharmaceutical to test in future human clinical trials (https://www.prnewswire.com/news-releases/biomarin-highlights-breadth-of-innovative-development-pipeline-at-rd-day-on-october-18th-in-new-york-300539018.html).

While loss of frataxin is believed to be the main driver of the disease, the complex pathophysiology of FRDA is still not fully elucidated. For example, the roles of oxidative stress and iron metabolism in FRDA pathology are unclear (21,22). Additionally, similar to other neurodegenerative diseases, only certain cell types and tissues are affected, despite frataxin being ubiquitously expressed. Previous gene expression profiling studies aimed at addressing the molecular basis of FRDA pathophysiology have been conducted in mouse models that do not to fully recapitulate the human disease (18,23-27) or human cells that do not represent an affected tissue (28-34). The advent of induced pluripotent stem cell (iPSC) technology (35) has allowed in vitro modeling of diseases that involve inaccessible human tissue (36). Moreover, advances in genome editing techniques are allowing the establishment of isogenic lines that overcome inter-individual variabilities in genome-wide studies. Here we present the first transcriptomic study in FRDA of human iPSC-derived central nervous system (CNS) and isogenic sensory neurons (SNs) and identify distinct but linked dysregulated pathways that are partially restored by HDACi treatment.

## Results

### Transcriptional profiling of FRDA iPSC-derived neuronal cells

We have previously derived iPSC lines from FRDA fibroblasts (37) and shown that they can be differentiated into functional β-III tubulin-positive neurons (20). Using a modified version of our published protocol (adapted from (38)), we differentiated four iPSC lines, two from unaffected individuals (KiPS, (39) and GM08333, Coriell Institute) and two from FRDA patients (from Coriell fibroblast lines GM03816 and GM04078) into CNS neurons. To investigate the effect of loss of frataxin on global gene expression and the effect of HDACi **109** (20) on such transcriptional changes, 14 day-old neurons were treated for 48 hours with 5 µM **109** or DMSO. *FXN* expression is lower in FRDA neurons compared to controls and is increased upon **109** treatment in affected cells (Fig. 1A). Total RNA was extracted and rRNA-depleted RNA samples were sequenced using the Illumina HISeq Analyzer 2000 and mapped to the human genome (see Experimental Procedures). Over 10 million single-end 75 bp reads were generated from each sample, with ~70-75% of the reads mapping to exons. Principal component analysis (PCA) shows clustering dependent on *FXN* expression and HDACi treatment (Fig. 1B). 5545 genes were found to be differentially expressed (DE) between FRDA lines and controls (with a False Discovery Rate (FDR) < 0.01), of which 2939 were upregulated and 2606 were downregulated (Figs 1C and 1D). Remarkably, and as reported previously (18,30), over 50% of dysregulated genes are reverted towards normalization when patient cells are treated for 48h with HDACi **109** and about 10% of these changes are statistically significant (Fig. 1E). In order to investigate the nature of these DE genes and the effect of HDACi **109** on their expression, we performed weighted gene coexpression network analysis (WGCNA) to identify modules of coexpressed genes that share similar functions or cellular localization. Dendrograms of clustered genes, identified modules and their correlation are shown in Fig. 2A and 2B and module membership for each gene is indicated in supplementary file 1. Based on an FDR cutoff of <0.1 and on the module eigengenes (Fig. 2C), which summarize each module expression data across all samples, six modules were selected for pathway analysis. The brown, magenta and black modules include genes that are downregulated as a consequence of repeat expansion, while the pink, turquoise and yellow modules include genes that are upregulated in the disease state. In the magenta and pink modules, the expression of dysregulated transcripts changed towards normalization with **109** treatment, which is possibly a consequence of restoration of *FXN* transcription. Top enriched pathways identified within the GO biological processes and cellular component databases are shown in Fig. 2D. It is notable that overrepresented genes in the pink module are related to transcriptional regulation and that these genes are reverted toward normalization by treatment with the HDAC inhibitor. Other identified pathways are extracellular matrix (ECM) organization (brown), chemical synapsis transmission (turquoise), regulated exocytosis (magenta), mitochondrion (black) and vesicle-mediated transport (yellow). While the black module points to the well-established mitochondrial dysfunction in FRDA, we failed to see a beneficial effect of **109** on the expression of these genes.

**Figure 1.**
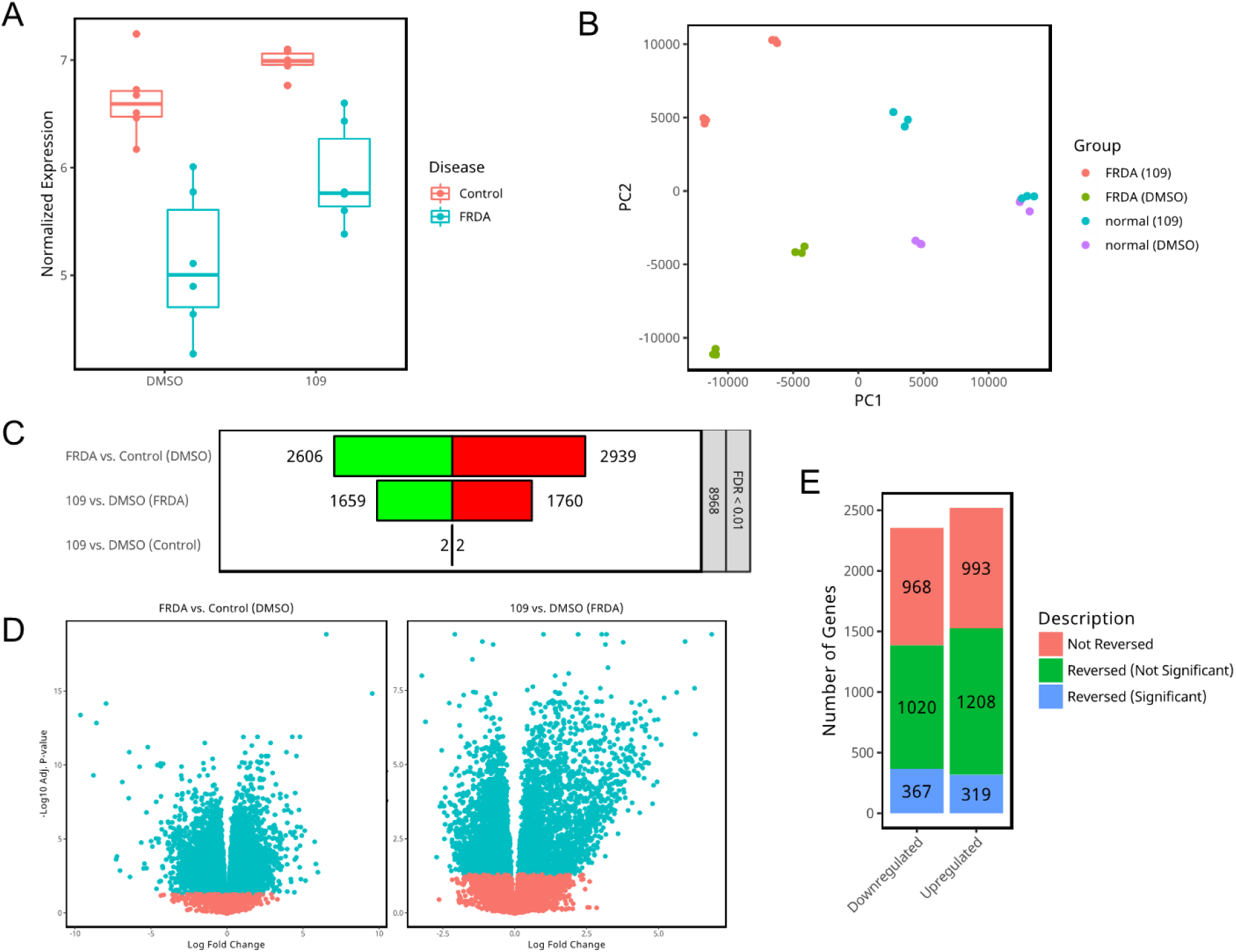
Transcriptomic analysis of FRDA and unaffected iPSC-derived neurons. *A*, Frataxin expression in two FRDA and two control neuronal lines, plotted as voom-transformed values from RNA-seq counts. *B*, PCA plot of samples. *C*, Number of differentially expressed genes with FDR < 0.01, for the comparisons shown. Green (left panels) are downregulated and red (right panels) are upregulated genes. *D*, Volcano plots of DE genes in FRDA versus control cells (left) and 109-treated versus DMSO-treated FRDA neurons. The x-axis shows the log fold change (logFC) and the y-axis show the -log10 p-values for each gene. Points in green represent genes that are significant at FDR <0.01 while points in red represent genes that were not significant. *E*, Changes in DE genes between FRDA and control neurons upon HDACi 109 treatment.

**Figure 2.**
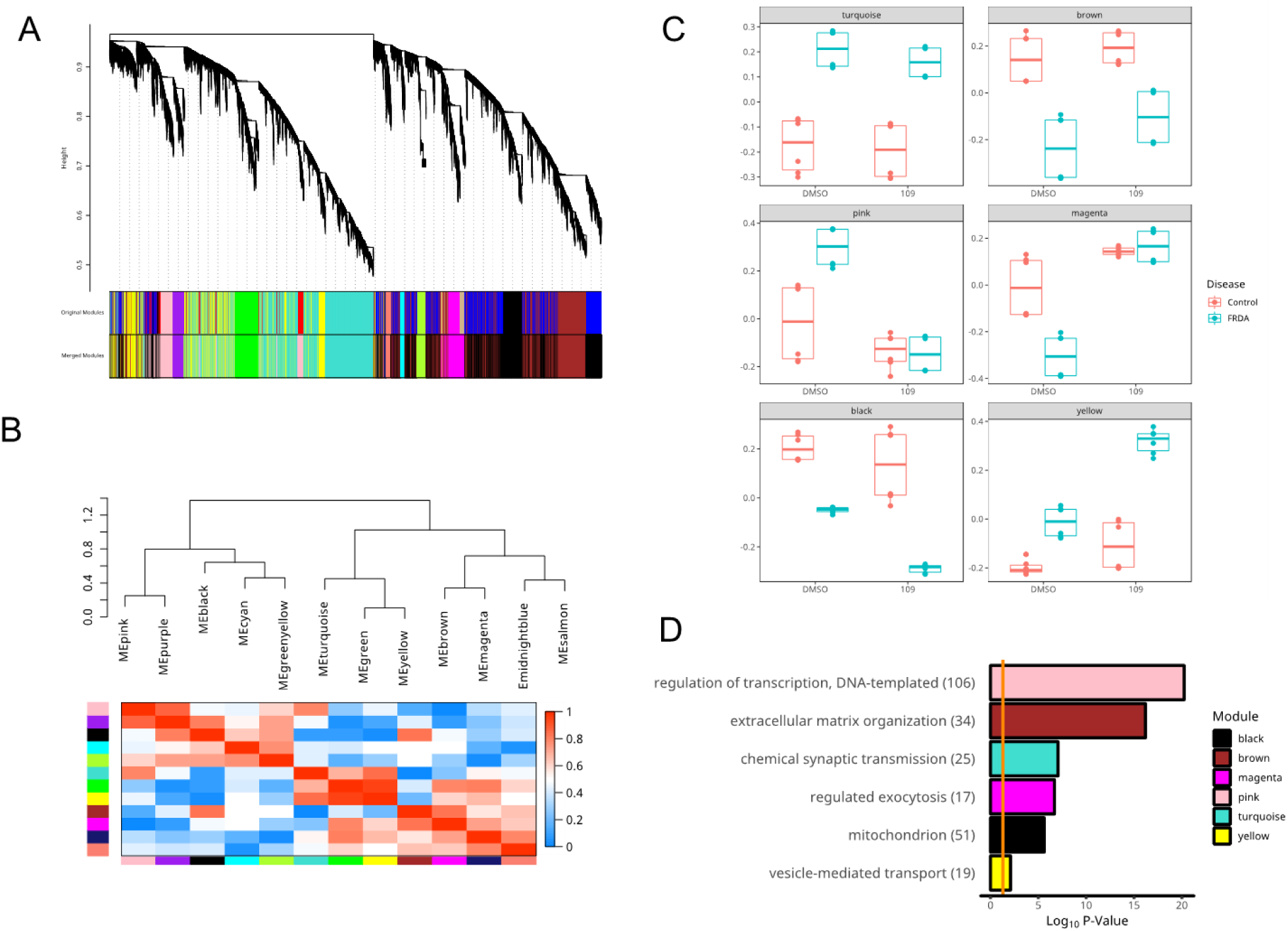
Network analysis. *A*, Hierarchical clustering dendrogram of all genes in the co-expression network. Module assignments are shown in the bar at the bottom. *B*, Top: Hierarchical clustering dendrogram of module eigengenes. Bottom: Heatmap of correlation coefficients between each pair of module eigengenes. *C*, Eigengenes for the 6 most significant pathways (FDR<0.1) *D*, Top overrepresented pathways in the six selected WGCNA modules.

### Creation of isogenic iPSC lines by adenovirus-mediated homologous recombination

The comparison of transcriptomes of unaffected and diseased iPSC-derived neuronal cells can be confounded by their different genetic backgrounds (40). We therefore sought to derive isogenic iPSC lines that differ exclusively in the number of GAA•TTC repeats in the first intron of the *FXN* gene. Compared to nuclease-based methods such as zinc finger nucleases, TALENs and CRISPR-Cas9 systems, homologous recombination offers the advantage of virtually scar-less gene editing. We selected helper-dependent adenovirus (HdAV) due to its capacity of accommodating large amounts of genetic material, absence of genomic integrations and its minimal mutation rate (41-44). We constructed a correction vector plasmid, *FXN*-*HdAV* that contained ~19 Kb of the human *FXN* gene with 6 GAA•TTC repeats in intron 1. The vector also included an excisable neomycin cassette flanked by *Frt* sequences that was used for selection and subsequently removed seamlessly by *Flp* recombinase excision (Supp. Fig. 1A).

Adenoviruses packaged with the linearized *FXN*-*HdAV* construct were used to infect FRDA patient-derived iPSCs, which we had previously generated and characterized (37) (from Coriell fibroblast line GM03816; hereby denoted as *FXN* ^exp/exp^). We then screened for clones that were corrected at one allele, using standard PCR methods. The removal of the neomycin cassette produced the isogenic heterozygous iPSC line *FXN*^exp/6^. By repeating this process, we then generated iPSC clones where both alleles were corrected (generating *FXN*^6/6^; see Supp. Fig. 1B and 1C for correction scheme). These isogenic iPSCs were characterized by karyotyping, embryoid body formation and pluripotency marker expression, confirming that they retained pluripotency (Supp. Fig. 2).

### Correction of the GAA•TTC repeats restores FXN expression and reverses heterochromatin formation in iPSCs and iPSC-derived neurons

The presence of short GAA•TTC repeats in the isogenic iPSCs was confirmed by standard PCR (Fig. 3A). *FXN* gene and frataxin protein expression were measured by quantitative real-time polymerase chain reaction (qRT-PCR) (Fig. 3B) and western blotting (Fig. 3C), respectively, demonstrating that correction of the GAA•TTC repeats restores *FXN* mRNA and protein levels to those of unaffected cells. To demonstrate reversal of heterochromatin silencing of the *FXN* gene, chromatin immunoprecipitation (ChIP) studies were performed and revealed that H3K9 acetylation, a critical histone mark downregulated in FRDA cells (20), was restored to normal levels in fully corrected *FXN*^6/6^ iPSCs (Fig. 3D). The expression of the *FXN* transcript and of frataxin upon repeat shortening clearly demonstrates that the repeat expansion is solely responsible for the transcriptional defect. Moreover, the acetylation of histones around the GAA•TTC repeats is directly correlated to repeat length.

**Figure 3.**
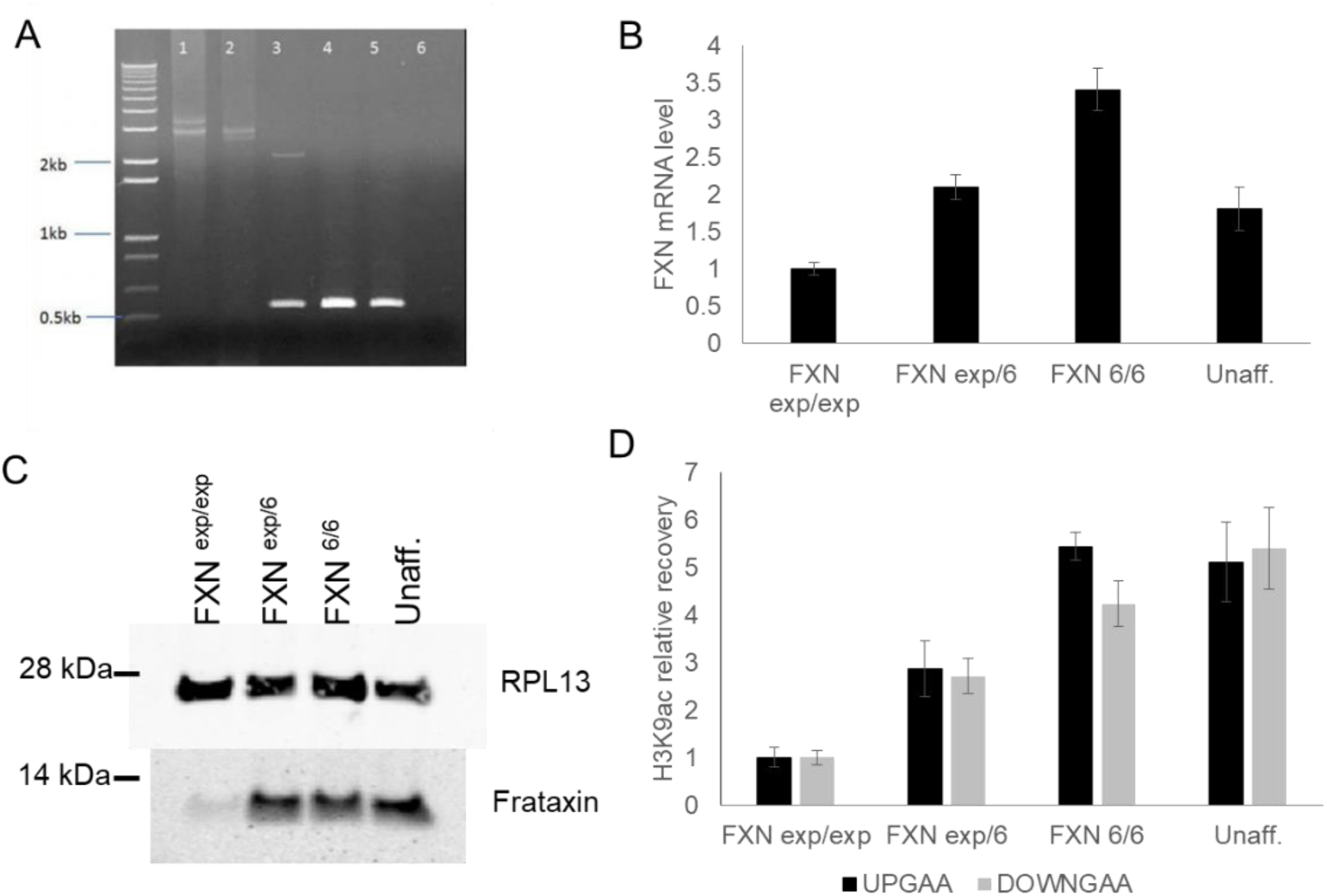
Characterization of isogenic iPSC lines. *A*, Genotyping of lines FXN^exp/exp^ (1), FXN^exp/6^ before neomycin cassette removal (2), FXN^exp/6^ after neomycin cassette removal (3), FXN^6/6^ (4) and one non-isogenic unaffected control (5). Lane 6 is a no DNA negative control. Ethidium bromide stained agarose gel is shown. *B*, qRT-PCR quantification of *FXN* mRNA levels in the isogenic lines and in one unaffected non-isogenic line (WT). *C*, Western blot of cell lysates derived from isogenic iPSCs and one non-isogenic control, probed with anti-frataxin and anti-RPL13 antibodies. The position of molecular weight markers is indicated to the left *D*, Levels of H3K9 acetylation as measured by ChIP upstream (up GAA) and downstream (down GAA) of the GAA•TTC repeats in the same iPSC lines as in *B*.

The three isogenic iPSC lines, *FXN*^exp/exp^, *FXN*^exp/6^ and *FXN*^6/6^ were differentiated by dual SMAD inhibition (38,45) into neuronal cells that expressed β-III tubulin (Supp Fig. 3A) and other neuronal markers such as *HUC* and *MAP2* (as measured by qRT-PCR, Supp Fig. 3B).

Similarly to the isogenic iPSCs, the isogenic corrected neurons showed a restoration of *FXN* gene expression when compared to FRDA neurons (Supp. Fig. 3C) and absence of repressive chromatin marks near the repeats (Supp. Fig 3D). The analysis of eight histone postsynthetic modifications (20) by ChIP at the *FXN* promoter and in the regions upstream and downstream of the GAA•TTC repeats shows increased acetylation and decreased methylation in the corrected *FXN*^6/6^ neurons compared to the isogenic *FXN*^exp/exp^ FRDA neurons. (Supp. Fig. 3D). Transcriptomic analysis of these isogenic β-III tubulin-positive cells was performed (not shown) and has been deposited in the Sequence Read Archive (SRA) (accession PRJNA495860).

### Transcriptional profiling of FRDA iPSC-derived sensory neurons

Signature neuropathological elements of FRDA are the thinning of dorsal root fibers of the spinal cord due to progressive loss of large DRG sensory neurons and low myelination of peripheral sensory nerves (5). We therefore sought to derive sensory neurons (SNs) from FRDA and isogenic control iPSCs to perform transcriptional profiling by RNA-sequencing. We adapted published protocols (46,47) that involve the forced expression of two transcription factors BRN3A and NGN1 and the use of small molecules that have been shown to induce the differentiation of sensory neurons of the nociceptor type. The combination of the two protocols (see Experimental Procedures) produced a more efficient differentiation than the two individual methods, as measured by the expression of the sensory neuron marker ISL1 (data not shown). Immunostaining analysis at day 11 after differentiation showed that these cells express β-III tubulin and peripherin (Fig. 4A). We also detected similar levels of expression of the SN marker ISL1 and NTRK3 (TRKC), together with PUO4F1 (BRN3A), in both the *FXN*^exp/exp^ and *FXN*^6/6^ lines, as measured by qRT-PCR (Fig. 4B). Transcriptomic analysis was performed as described above to compare 11-day-old sensory neurons derived from the *FXN*^exp/exp^ and *FXN*^6/6^ lines and their corresponding iPSCs. PCA plot of these samples shows clustering dependent on cell identity and *FXN* expression (Figs. 5A and 5B). When comparing iPSCs to sensory neurons from both patient and its isogenic control, over 12,000 DE genes for each comparison were identified (Fig. 5C), denoting the expected major changes in the trascriptome upon differentiation. The expression of a panel of sensory neuron markers in iPSCs versus SNs, as measured by RNA-seq voom-transformed values, confirms the identity of these cells (Fig. 4D). Nav1.7 (*SCN9A*), *TRPV1* and *P2RX3*, some of the most selectively expressed ion channels in human DRG (48) were upregulated in these neurons, together with *SLC17A7* and *SLC17A6* (glutamate transporters VGLUT1 and VGLUT2), while markers of fetal brain (SATB2 and POU3F2), astrocytes (GFAP), cerebellum (EOMES), melanocytes (MITF) and Schwann cells (MPZ) were not highly expressed (Fig. 4D).

**Figure 4.**
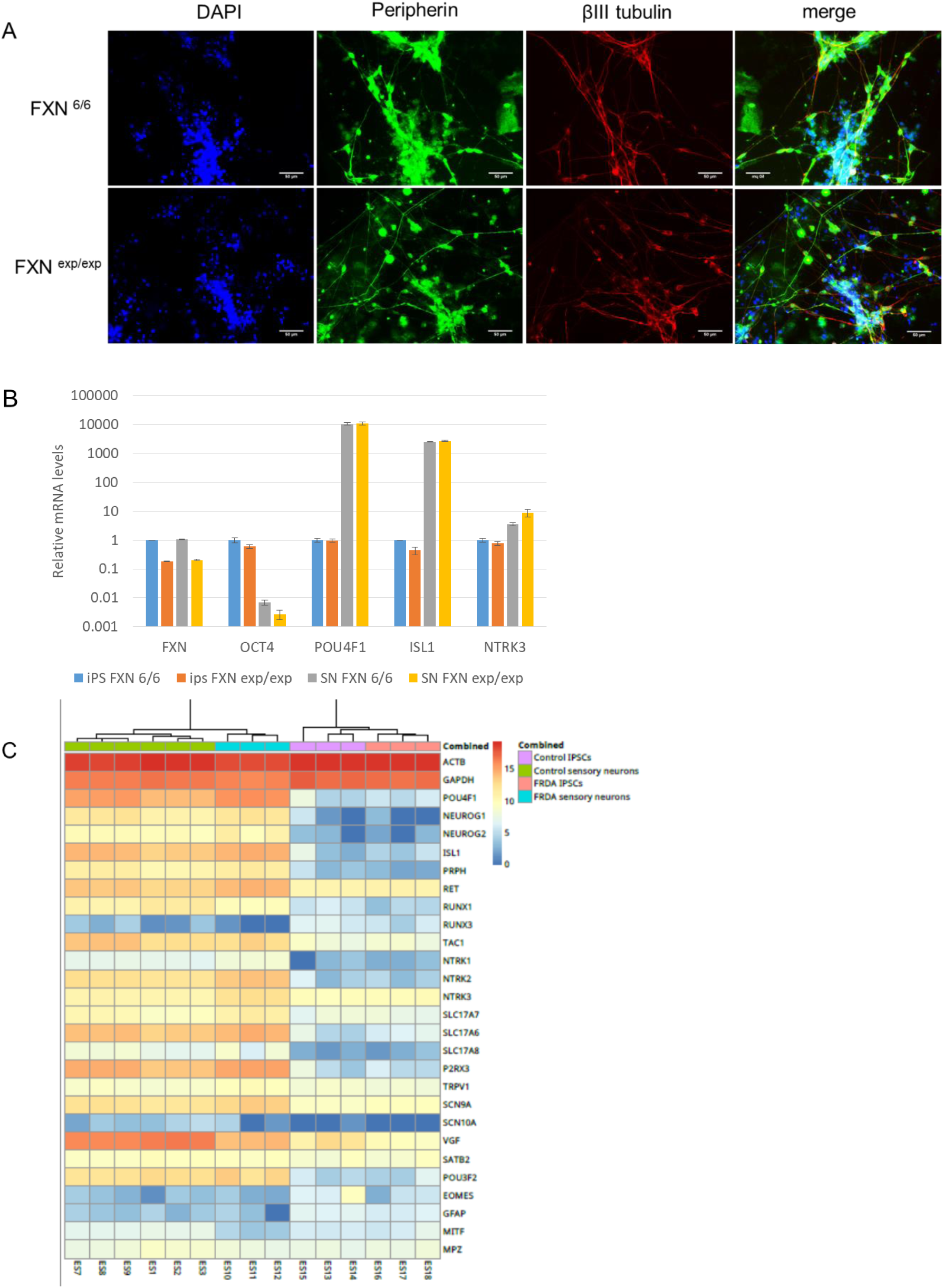
iPSC-derived sensory neuron characterization. *A*, Immunofluorescence staining of 11-day-old sensory neurons with the pan neuronal marker β-III tubulin (red) and the peripheral nervous system marker peripherin (green). Nuclei were counterstained with DAPI. Scale bar = 50 µM. *B*, Expression of FXN, the pluripotency marker OCT4 and sensory neuron markers POU4F1, ISL1 and NTRK3 as measured by qRT-PCR in iPSCs and SNs. *C*, Heatmap showing expression of a panel of sensory neurons markers in isogenic iPSC lines (both FRDA and control, right) and isogenic SNs (both FRDA and control, left). Shown is also the expression of two housekeeping genes (ACTB and GAPDH) together with markers of other tissues and cell types: SATB2 and POU3F2 (fetal brain), EOMES (cerebellum), GFAP (astocytes), MITF (melanocytes), MPZ (Schwann cells).

**Figure 5.**
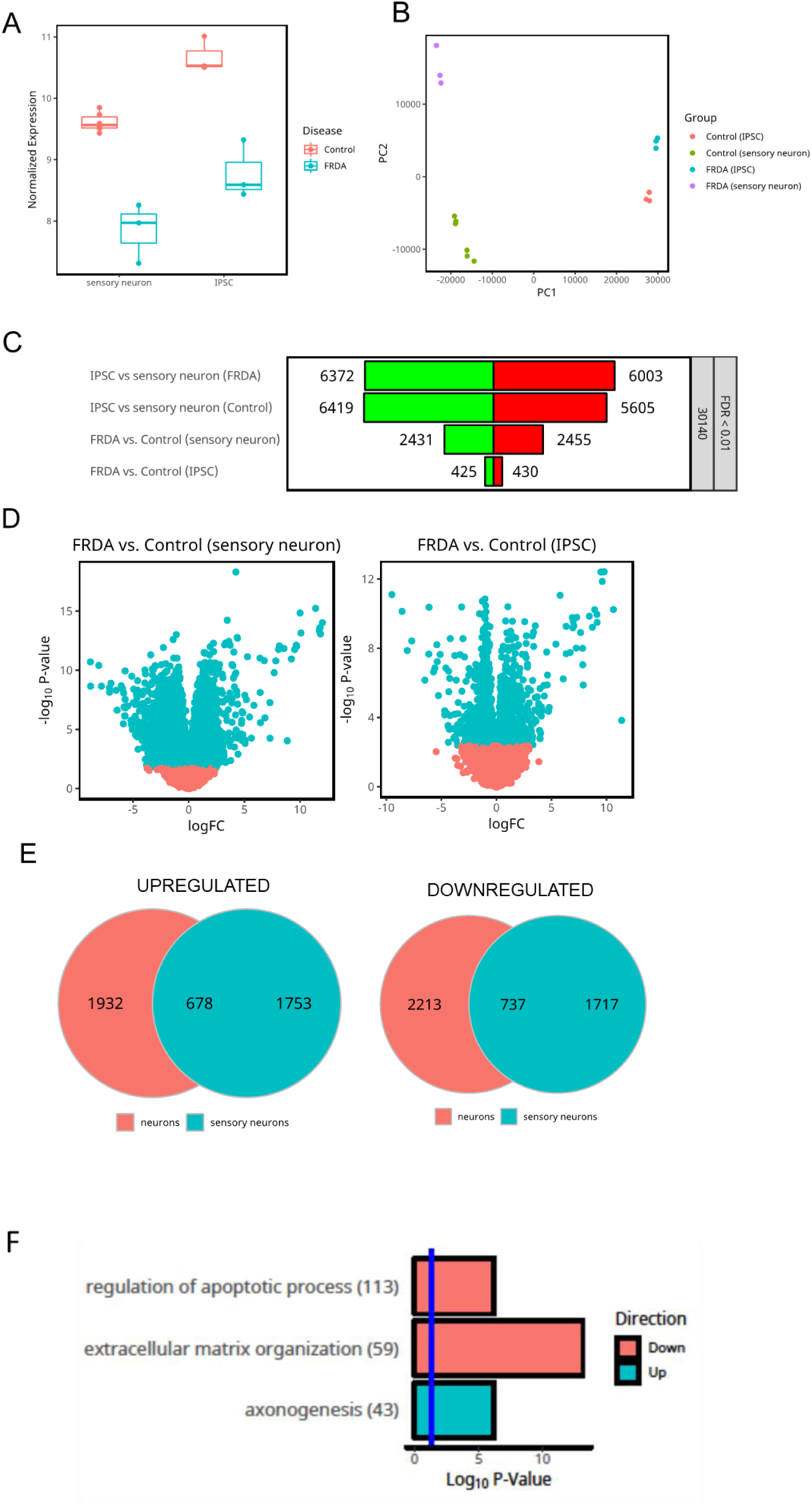
Transcriptomic analysis of FRDA and unaffected iPSC-derived sensory neurons. *A*, Frataxin expression in isogenic iPSCs and SNs, plotted as voom-transformed values from gene counts. *B*, PCA plot of samples. *C*, Number of differentially expressed genes with FDR < 0.01, for the comparisons shown. Green (left panels) are downregulated and red (right panels) are upregulated genes. *D*, Volcano plots of DE genes in FRDA versus control SNs (left) and FRDA versus control iPSCs. The x-axis shows the log fold change (logFC) and the y-axis show the -log10 p-values for each gene. Points in green represent genes that are significant at FDR <0.01 while points in red represent genes that were not significant. *E*, Venn diagrams of upregulated (left) and downregulated (right) DE genes common to CNS and PNS neurons. *F*, Pathway analysis of shared DE genes between CNS and PNS neurons.

Notably, when comparing FRDA and unaffected cells, only 855 genes differed in iPSCs while 4886 were differentially expressed in SNs (Figs. 5C and 5D). This could represent an expansion of the FRDA signature in a relevant cell type, but also be the result of differentiation-induced variability in the two lines, as reported in a recent study (40). We identified 2431 genes that were downregulated and 2455 genes that were upregulated in *FXN*^exp/exp^ SNs compared to their isogenic controls and among these, 678 and 737 genes respectively, were common to CNS neurons (Fig. 5E). Interestingly, the common downregulated genes are uniquely enriched for regulation of apoptosis pathways, while ECM organization-related terms were also identified within the brown module (Fig. 2D and 5F). Upregulated genes that are common to CNS and peripheral nervous system (PNS) neurons are overrepresented in genes for axonogenesis (Fig 5F).

### A gene expression signature of FRDA

Our RNA sequencing studies of iPSC-derived neurons provide evidence of metabolic changes in FRDA cells versus controls. These changes are small but involve multiple aspects of cell metabolism. FRDA neurons show a decrease in mitochondrial protein transcript levels, such as components of the ATP synthase complex and of complex I (black module), all of which have been described in FRDA (23,27,49-51). We observe consistent changes in ECM organization, focal adhesion and related signaling (brown and magenta modules and SNs), possibly linked to cytoskeletal abnormalities that have been reported in FRDA cells (52-54). We also identify changes in chemical synapsis transmission (turquoise module, Fig. 1D) and axonogenesis (SNs). Lastly, we detect widespread changes in the regulation of transcription (pink module). This is very intriguing because metabolism and signaling events are intimately entwined and recent evidence indicates that the two regulate each other (55-57). Remarkably, the comparison of DE genes in CNS and sensory neurons provides evidence of involvement of apoptotic processes. Taken together, these transcriptional changes represent a gene expression signature of frataxin loss and, possibly, of the disease.

## Discussion

It is widely accepted that the loss of frataxin is the causative event of FRDA pathophysiology, albeit other genetic modifiers of disease severity could exist. Despite this apparent simple etiology, the mechanisms of disease pathogenesis remain unclear. This is in part due to the fact that although frataxin has been shown to be an essential component of Fe-S cluster assembly (reviewed in (6)), its function is not yet fully understood. Defects in Fe-S proteins like aconitase and respiratory chain complex components, together with oxidative stress induced by iron overload, defects in lipid metabolism, cytoskeleton assembly and heme biosynthesis have been implicated in FRDA pathophysiology (21,22,58,59). The aim of this study was to further disease understanding, by providing transcriptome analysis of a relevant affected tissue in FRDA. The compelling hypothesis that the tissue specificity of human disease is linked not simply to the expression of a particular gene, but to a “module” or subnetwork of genes that all have to be expressed in the same tissue in order for the disease to manifest (60), renders transcriptional profiling studies essential to understand the tissue specificity of FRDA.

Examples of studies employing iPSC-derived disease-relevant cells are ample in the literature (reviewed in (36,61-63)), but the use of these models also uncovers numerous pitfalls such as the lack of maturity or sign of aging in these cells (64-67) and the variability introduced by genetic background and differentiation (40,68). We used relatively “young” or immature cells, focusing on a recent report that implicates hypoplasia rather than atrophy in the demise of the DRG(5). According to this hypothesis, “young” and immature iPSC-derived neurons could provide information on early disease mechanisms. Our FRDA CNS neurons do not show any phenotypical deficit like the ones described for other FRDA cell types or animal models (27,50,51,53,69-78). Complex I, Complex III and aconitase activity were all similar to unaffected neurons and so were oxygen consumption, spare respiratory capacity, ATP production, reactive oxygen species formation and mitochondrial membrane potential (MMP, data not shown), while others have shown reduction in MMP(79), increased oxidative stress and decreased levels of Fe-S cluster- and lipoic acid-containing proteins (80) in iPSC-derived FRDA neurons compared to controls. We therefore expected the transcriptional changes to be subtle. To distinguish between disease-related changes and noise introduced by line-to-line variability during differentiation, we treated both unaffected and FRDA neurons with the HDACi **109**, thus restoring *FXN* transcription in the FRDA neurons (Fig. 1A and (20)). We reasoned that changes that are a true consequence of the disease state would also be reverted by treatment with **109**. In fact, over 50% of DE genes were reverted toward normalization by **109** treatment, similarly to what we have shown previously (18,30). WGCNA also allowed us to focus pathway analysis on modules of genes that shared similar expression profiles. This led to cleaner and more interpretable gene set annotation, as genes in the same pathway tend to show highly correlated gene expression patterns.

As noted above, our RNA sequencing data revealed subtle changes in expression of genes involved in cell metabolism. Changes in metabolism, like glucose utilization, can translate to changes in protein posttranslational modifications, such as glycosylation and acetylation, and these modifications are sensors in the cell of nutrient availability (55). For example acetylCoA is highly compartmentalized and although its levels are not limiting in the cell, fluctuations in local availability can result in changes in protein acetylation and epigenetic alterations (81) and can account for changes in the regulation of transcription identified in the pink module (Fig. 2D). It is notable that hyperacetylation of mitochondrial proteins occurs in the heart of a conditional *Fxn*-knockout mouse model of FRDA (82,83). Changes in transcription regulatory activity were also detected in the heart and cerebellum of an inducible mouse model of frataxin deficiency (FRDAkd mouse (27)). As noted above, we also identified pathways related to chemical synapsis transmission (turquoise module, Fig. 1D). Upregulation of synapsis related genes were also reported in the FRDAkd mouse (27). Axonogenesis was the most enriched term for upregulated genes shared between CNS and sensory neurons. This term was also enriched in the turquoise module, but not as significantly (not shown). Whether this upregulation is a compensatory event (for examples because of defects in ECM-cytoskeleton interactions that could cause axonal retraction) or a result of variability induced by differentiation (40) remains to be determined.

As shown above, the expression of more than half of DE genes between FRDA and unaffected neurons changes towards normalization upon **109** treatment, although only about 10% of these changes are statistically significant. While we detected 3419 DE genes when comparing DMSO and HDACi **109**-treated FRDA cells (with FDR>0.01), only 4 genes were significantly different in unaffected cells (fig 1C). This is unlike previous results, where we showed that hundreds of genes were changed upon HDACi **106** treatment (a reverse amide of HDACi **109** (84)) in unaffected lymphocytes(30), and probably due to line-to-line variability in the response to HDACi (see PCA plot in 1B), as well as the biological differences between iPSC-derived neurons and blood cells. Among the most significant changes upon **109** addition were genes involved in regulated exocytosis (magenta module) and regulation of transcription (pink module), but not mitochondrial proteins (black module). It is conceivable that a more prolonged **109** treatment and sustained increase in frataxin protein are necessary to see more extensive correction of the transcriptional changes.

To address the concern of line-to-line variability due the different genetic backgrounds, we created isogenic iPSC lines. We elected to utilize homologous recombination as opposed to nuclease-based methods (such as ZFNs, TALENs and CRISPR-Cas9) because of the nature of the *FXN* gene near the GAA•TTC repeats. Since the repeats are embedded in an Alu element (85), selection of nuclease sites are problematic. Moreover, it is not known whether the sequence flanking the repeat might be important for *FXN* gene expression or RNA processing. For example, epigenetic indicators of regulatory regions like H3K27ac peaks are detectable upstream of the repeats (S. Bidichandani, personal communication). We selected HdAV because of its large capacity to incorporate genetic correction material (up to 30 kilobases) and no possibility of nuclease mediated off-targeted cleavage (41-44). This methodology utilizes a “helper” virus that aids in packaging the desired vector into replication deficient adenoviruses. The packaged adenovirus possesses no ability of replication, but retains the ability of infection and highly efficient delivery of its genetic content. We used two rounds of homologous recombination to correct both alleles in *FXN*^exp/exp^. The resulting *FXN*^6/6^ iPSC line showed a restoration of *FXN* transcription and histone acetylation around the GAA•TCC repeats, comparable to that of unaffected cells. This unequivocally demonstrates that the repeat expansion is solely responsible for the transcriptional defect and that acetylation of histones in this region is determined by repeat length.

Given that the peripheral nervous system, and in particular the DRG, are the initial site of neurodegeneration, it was important to perform additional transcriptional profiling by RNA-seq using the most relevant cellular model, noting that β-III tubulin positive neurons differentiated by dual SMAD inhibition represent a model of the central nervous system (45). We therefore derived sensory neurons (SNs) from FRDA and isogenic control iPSCs by adapting protocols from the Baldwin (46) and Studer (47) groups that involve the forced expression of two transcription factors, BRN3A and NGN1, and the use of small molecules that have been shown to induce the differentiation of sensory neurons of the nociceptor type. The sensory neuron identity of these cells is demonstrated by overexpression of SN markers like *ISL1* and *POU4F1* (BRN3A) mRNAs, detected both by qRT-PCR and RNA-seq (Fig. 4). Receptors specific to SN subtypes (TRKA-C) are all overexpressed compared to iPSCs, although we could not detect the expression of RUNX3, which controls the development of proprioceptive, TRKC-positive neurons (86). Transcriptomic analysis showed that other markers of sensory neurons are expressed in these neurons like Na_v_1.7 (*SCN9A*), *TRPV1* and *P2RX3* (Fig. 4D). These genes encode some of the most selectively expressed ion channels in the human DRG (48). Markers of fetal brain (SATB2 and POU3F2), astocytes (GFAP), cerebellum (EOMES), melanocytes (MITF), Schwann cells (MPZ) are either very poorly expressed or unchanged during SN differentiation.

Despite the creation of isogenic lines being expected to reduce differences associated with diverse genetic backgrounds, we anticipated that we would still detect some level of differentiation variability in the two isogenic lines.

Almost 30% of the DE genes in sensory neurons were common to CNS neurons and represent some of the same pathways that were discussed above. These common genes, however, were uniquely enriched for genes involved in the regulation of apoptosis. Involvement of apoptotic mechanisms has been previously reported in FRDA cells (53,70,72,87,88). This suggests that to overcome the pitfalls of studies centered on iPSC-derived cell models, both multiple cell lines and isogenic lines are necessary. Genes that are specifically dysregulated in SNs but not CNS neurons could represent a true disease signature rather than the result of changes that compensate for frataxin deficit in cell types that do not appear to be affected in the disease. Pathway analysis of these genes showed enrichments very similar to the genes in the WGCNA modules. This could indicate that, because of the immature nature of these cells, the same transcriptional changes are occurring in the two neuronal types. The question remains whether and how any of the identified changes in gene expression contribute to pathogenesis. All or some could be compensatory mechanisms of frataxin deficiency and an adaptive response to metabolic changes (since no overt phenotype was detectable). It is conceivable that, at some point during development or aging, this adaptive response could become ablated and the cell overwhelmed by frataxin loss, and neurodegeneration occurs. We hypothesize that failure to observe an FRDA phenotype in iPSC-derived neuronal cells could be due to the immature nature of such cells, and the fact that most FRDA patients present with symptoms between the ages of 8 to 15 years, with no symptoms at birth. Since generation of iPSCs from patient fibroblasts resets the developmental clock back to the embryonic stem cell stage, it is not that surprising that iPSC-derived cells fail to recapitulate the degenerative hallmarks of FRDA. Several approaches are currently available to generate “aged” neurons from either donor fibroblasts (89) or forced aging of iPSC-derived cells by expression of progerin(64). Future studies in FRDA disease modeling could use these methods to attempt to recapitulate an FRDA phenotype in vitro.

## Experimental Procedures

### Neuronal differentiation

iPSC line generation and characterization was described previously(37). Neuronal differentiation was performed as described (38) with minor modifications. Briefly, iPSCs grown on Matrigel (Corning, Coming, NY) and mTeSR (Stem Cell Technologies, Vancouver, Canada) were treated for 10 days in E6 medium (Stem Cell Technologies, Vancouver, Canada) with 0.5μM LDN-193189, 10μM SB431542 and 20μg/ml FGF2. iPSC colonies were dissociated with Accutase (Innovative Cell Technologies, San Diego, CA) and plated in AggreWell plates (Stem Cell Technologies, Vancouver, Canada) at a density of 1000 cells per microwell. Neurospheres were grown in suspension for one week in Neurobasal A medium (Thermo Fisher Scientific, Waltham, MA), supplemented with N2 and B27 supplements (Thermo Fisher Scientific, Waltham, MA) and FGF2 and EGF, both at 20μg/ml. Neurospheres were then plated on Matrigel and grown in the same medium as above until rosettes appeared. Rosettes were manually isolated, grown for 4 to 7 days in suspension, then dissociated with Accutase and plated on Matrigel at 200,000 cells/cm^2^. To induce neuronal differentiation, cells were grown in the same medium as above without FGF2 and EGF for 14 days.

For sensory neuron differentiation, iPSCs were plated at 50,000 cells/cm^2^ in mTeSR and infected with lentiviruses expressing a bicistronic construct of *POU4F1* (Brn3a) and *Neurog1* (Ngn1) under the control of a doxycycline inducible promoter and the transactivator rtTA (90)(see plasmid section below). The next day (day 1) medium was changed to mTeSR supplemented with 5μg/ml doxycycline. On day 2 medium was changed to N2 medium (46) with 0.5μM LDN-193189, 10μM SB431542 and 5μg/ml doxycycline. On day 4 medium was changed to N2 medium with 0.5μM LDN-193189, 10μM SB431542, 5μM CHIR99021, 10μM SU5402, 10μM DAPT and 5μg/ml doxycycline. On day 5 medium was changed to N3 medium(46) with 5μM CHIR99021, 10μM SU5402, 10μM DAPT and 5μg/ml doxycycline. On day 7 medium was changed to N3 medium with 10μM SU5402 and 10μM DAPT. On day 10 medium was changed to Neurobasal A supplemented with N2 and B27. Cells were collected at day 12 for total RNA isolation.

### RNA sequencing, differential expression and WGCNA

RNA was isolated using the RNeasy mini kit (QIAgen, Hilden, Gemany). Single end 75bp reads were generated by the NextSeq (Illumina, San Diego, CA) located at the Scripps Next generation Sequencing Facility. Image analysis, base calling, and demultiplexing was done using bcl2fastq. Cutadapt is used to trim the adapter and low base pair called scores. FASTQ files were aligned to BAMs and counts were generated using STAR (91). Raw counts were transformed to counts per million using limma voom (92) and were normalized for library size using TMM normalization implemented by edgeR (93). Differential expression was analyzed using linear models with limma (94), and no additional covariates were added to the models. Network analysis was run with the WGCNA package (95), using a soft power threshold of 7 for the adjacency matrix, and a minimum module size of 20. Modules with a correlation higher than 0.8 were merged. Gene sets analyzed for enrichment analysis were submitted to Enrichr (96,97), with pathways being identified in the GO Biological Process database, and cell compartments from GO Cellular Component.

### Plasmids

#### HdAV-FXN plasmid

A neomycin cassette containing the neomycin resistance gene flanked by Frt sequences under the control of bacterial and mammalian promoters was amplified from plasmid *pgk-gb-frt-neo-frt* together with 50 bp homology arms from the *FXN* gene, and inserted by recombineering (98) 108 bp downstream of the GAA•TTC repeats in the bacterial artificial chromosome (BAC) plasmid RP11-265B. 18.7 kb of this construct containing the human *FXN* gene locus with 6 GAA•TTC repeats and the neomycin cassette was cloned in pGEX-4T1 by recombinnering(98). To construct the HdAV-FXN plasmid, the 18.7 kb fragment was isolated by restriction digestion and subcloned in the adenoviral plasmid pCIHDAdGT8-3.

#### Bicistronic plasmid expressing Brn3a and Ngn1

Mouse Neurog1 cDNA (83% homologous to human cDNA peptide) was cloned without the stop codon and replaced with an F2A self-cleaving peptide sequence. Human BRN3A cDNA (*POU4F1*) was added directly after the F2A peptide to generate an Ngn1-F2A-Brn3a bicistronic sequence. The bicistronic sequence was inserted into a lentiviral construct under the control of the tetracycline operator (TetO) as described previously(99). Replication-incompetent VSVg-coated lentiviral particles were packed in HEK293T cells (ATCC), collected 48h after transfection, and filtered through a 45μm membrane before use.

### HdAV virus particle production

Helper virus particle were produced by infecting HEK293T cells (Multiplicity of infection of 3). After 48 to 72 hours of incubation when cytopathic effect (CPE) was greater than 90% the cells were lysed by 3 freeze-thaw cycles and the virus purified by cesium chloride gradient centrifugation (100). The HdAV particles were assembled according to previously published methods (101). Briefly the HdAV-FXN plasmid was transfected into 116 cells (a modified HEK293 cell line that expresses high levels of Cre recombinase(102)) and these cells subsequently infected with the purified helper virus at a MOI of 0.2-0.3. when 90% CPE was detected, the crude cell lysate was collected and use to re-infected 116 cells. Multiple rounds of infections were performed until the desired titer was achieved as measured by β-galactosidase assay. The cell lysate was purified and concentrated by CsCl centrifugation (100).

### Polymerase Chain Reaction (PCR), quantitative Reverse Transcriptase Polymerase Chain Reaction (qRT-PCR), quantitative Western blotting and chromatin immunoprecipitation (ChIP)

GAA•TTC repeat length determination was performed as previously described (103). qRT-PCR and quantitative western blotting were previously described (37). Primer sequences for mRNA quantification were as follows: for *FXN* transcript, 5’-CAGAGGAAACGCTGGACTCT-3’ and 5’-AGCCAGATTTGCTTGTTTGG-3’, for POU4F1 transcript 5’-GGGCAAGAGCCATCCTTTCAA-3’ and 5’-CTGTTCATCGTGTGGTACGTG-3’, for ISL1 transcript 5’-GCGGAGTGTAATCAGTATTTGGA-3’ and 5’-GCATTTGATCCCGTACAACCT-3’, for NTRK3 transcript 5’-ACGAGAGGGTGACAATGCTG-3’ and 5’- CCAGTGACTATCCAGTCCACA-3’. Primer sequences for MAP2, ELAVL3 (HUC) and OCT4 were described in (37). The following antibodies were used in quantitative Western analysis: anti-frataxin (Santa Cruz Biotechnology, Dallas, TX) and anti-RPL13 (Cell Signaling Technologies, Danvers, MA), goat anti-mouse IR800 and goat anti-rabbit IR680, both from LI-COR Biosciences (Lincoln, NE). ChIP procedure and antibodies used are described in (20).

### Immunocytochemistry and FACS analysis

Immunocytochemistry and FACS analysis were described in (37). Primary antibodies included anti-β-III tubulin (Tuj1; Covance, Princeton, NJ), anti-Peripherin, anti-Oct4, anti-SSEA4, anti-Tra-1-60, (all from Millipore, Temecula, CA). Secondary antibodies were anti-mouse Alexa 594 and anti-rabbit Alexa 488 (Invitrogen, Carlsbad, CA).

## Acknowledgments

We thank Dr. Ulrich Müller for the gift of *pgk-gb-frt-neo-frt* plasmid and Dr. Steven Reed for the gift of pGEX-4T1. We also wish to thank Dr. Keiichiro Suzuki and Dr. Juan Carlos Izpisua Belmonte (Salk Institute, La Jolla, CA) for technical guidance and advice on the HdAV genetic correction method. The KiPS iPSC line was a gift from Dr. Jeanne Loring (Scripps). Research in the Gottesfeld lab is supported by grants from the National Institute for Neurological Disorders and Stroke (NINDS, 5R01 NS062856), the Friedreich’s Ataxia Research Alliance (grant to ES) and by BioMarin Pharmaceutical, Inc. J-IL was supported by a predoctoral fellowship from the Friedreich’s Ataxia Research Alliance (FARA). We acknowledge the support of the NINDS Informatics Center for Neurogenetics and Neurogenomics (P30 NS062691). DN was partially supported by the Friedreich’s Ataxia Research Alliance (FARA). Research in the Baldwin lab is supported by grants from the National Institute on Deafness and Other Communication Disorders (NIDCD, DC012592), the National Institute of Mental Health (NIMH, MH102698), the National Institute on Aging (NIA, DP1 AG055944), and the Dorris Neuroscience Center.

## Conflict of Interest Statement

JMG serves as a consultant to BioMarin Pharmaceutical for the development of histone deacetylase inhibitors as therapeutics, and is an inventor on patents licesence by The Scripps Research Institute to BioMarin Pharmaceutical. ES is supported in part by BioMarin Pharmaceutical.

The data discussed in this publication have been deposited in the Sequence Read Archive (SRA), accession PRJNA495860.

